# A simple approach for maximizing the overlap of phylogenetic and comparative data

**DOI:** 10.1101/024992

**Authors:** Matthew W. Pennell, Richard G. FitzJohn, William K. Cornwell

## Abstract

1. Biologists are increasingly using curated, public data sets to conduct phylogenetic comparative analyses. Unfortunately, there is often a mismatch between species for which there is phylogenetic data and those for which other data is available. As a result, researchers are commonly forced to either drop species from analyses entirely or else impute the missing data.
2. Here we outline a simple solution to increase the overlap while avoiding potential the biases introduced by imputing data. If some external topological or taxonomic information is available, this can be used to maximize the overlap between the data and the phylogeny. We develop an algorithm that replaces a species lacking data with a species that has data. This swap can be made because for those two species, all phylogenetic relationships are exactly equivalent.
3. We have implemented our method in a new R package phyndr, which will allow researchers to apply our algorithm to empirical data sets. It is relatively efficient such that taxon swaps can be quickly computed, even for large trees. To facilitate the use of taxonomic knowledge we created a separate data package `taxonlookup;` it contains a curated, versioned taxonomic lookup for land plants and is interoperable with phyndr.
4. Emerging online databases and statistical advances are making it possible for researchers to investigate evolutionary questions at unprecedented scales. However, in this effort species mismatch among data sources will increasingly be a problem; evolutionary informatics tools, such as phyndr and taxonlookup, can help alleviate this issue.

## Introduction

Phylogenetic comparative methods can be used to answer a broad range of evolutionary questions (O’Meara, 2012; Pennell & Harmon, 2013). At a practical level, doing so generally requires a phylogenetic tree and some set of species-level data; for example data on the species’ distribution, demography, species-interactions, physiology, or morphology. However, researchers commonly encounter a very mundane roadblock: some species have sufficient genetic data to build a phylogeny but have not been measured for traits of interest; others have been measured for the trait, yet are not placed within the phylogeny. To gain optimal power from comparative analysis, one generally wants to use as much data from both sources as possible, but the data mismatch prevents this.

This problem has become increasingly common: as the scale of phylogenetic comparative analyses expands — and fields outside of systematics find creative uses for phylogenetic data — researchers generally rely on previously published phylogenetic resources, in the form of sequence data and/or phylogenies, and trait data sets. There has been a recent push to assemble, curate, and open up, large collections of data, analogous to GenBank, for this purpose: TreeBASE (Sanderson *et al*., 1994) and Open Tree of Life (Hinchliff *et al*., 2015) for phylogenetic data and Encyclopedia of Life (Parr *et al*., 2014), try (Kattge *et al*., 2011), gbif (www.gbif.org), and compadre (Salguero-Gómez *et al*., 2015), among many others, for comparative data. There is also the common use of community presence/absence as a trait of interest (Vellend *et al*., 2011). The availability of phylogenetic data (both original sequence data and phylogenetic trees from published studies) is growing but is far from complete (Hinchliff & Smith, 2014), as is the case for traits. And both of these represent biased samples of life’s diversity — some groups of life and groups of traits have been studied much more intensely than others.

Consider the availability of data for vascular plants, a relatively well-studied group of organisms. There are 92,704 species for which there is currently *any* sequence data in GenBank (As of May 2015 — accessed using the ncbi taxonomy browser; Wheeler *et al*., 2007, http://www.ncbi.nlm.nih.gov/Taxonomy/Browser/ wwwtax.cgi). We compared this list to the 40,159 species included in a recent database of plant growth form (Zanne *et al*., 2014); the key limitation for comparative methods is the area of overlap between the two (Figure 1). While one dataset might be a strict superset of the other, in practice they contain overlap; we found 28,868 species represented in both data sets, with more species with trait data having genetic data than ther other way around.

**Figure 1:**
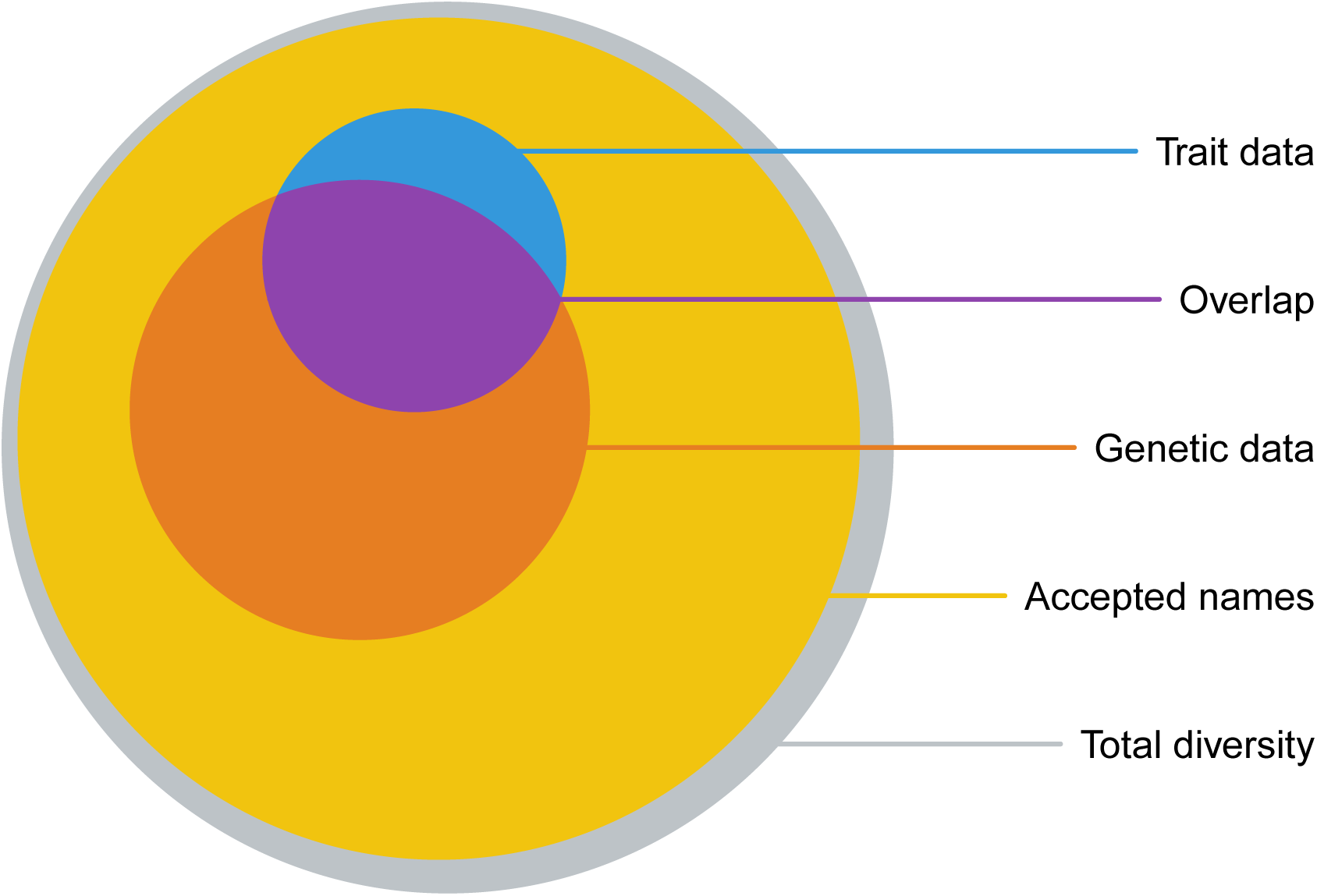
Overlap (purple) of species with recognized names (yellow), trait data in the global woodiness data base (blue Zanne *et al*., 2014; FitzJohn *et al*., 2014), and which have sequences deposited in GenBank (orange) (All, as of May 2015). The total diversity of plants (grey) is not known.

To increase the overlap (without gathering more data or estimating new phylogenies) a researcher is left with few options, all of which involve imputing data in some way. First, it is possible to add unplaced taxa into the phylogeny. If one is willing to assume the monophyly of some higher taxonomic group, it is possible to paste new terminal branches into the phylogenetic tree at approximately the correct location. However, neither the topological position nor the divergence time are known: one must either collapse the higher taxonomic group down to an (artificial) polytomy or randomly resolve relationships. Kuhn, Mooers & Thomas (2011) and Thomas *et al*. (2013) have suggested using a birth-death process, parameterized from the observed data to randomly resolve polytomies (see also Bapst, 2013, for a related approach for fossil trees) and this approach has been used to fill out trees for comparative analyses (Jetz *et al*., 2012; Price *et al*., 2012; Rolland *et al*., 2014; Jetz *et al*., 2014). For example, Jetz *et al*. (2012) produced a phylogeny containing all 9,993 species of birds but 3,323 (33.2%) of these lacked genetic data and were added in according to a constant rate birth-death process.

While such an approach may be very useful in some contexts, it may generate generate biases. A number of simulation studies have investigated this effect (Losos, 1994; Martins, 1996; Davies *et al*., 2012; Bapst, 2014; Rabosky, 2015) but the rationale is straightforward. If a unresolved clade in a rooted tree contains three taxa then the true phylogeny will only be sampled in 1/3 of random resolutions; more often than not, incorrect sister pairs will be generated. And if a trait of interest has any phylogenetic signal, then the sister species will be appear more divergent than they actually are, thus inflating the apparent rate of evolution. Of course, this problem quickly gets much worse as even more unplaced taxa are considered.

The problem of a mismatch between phylogenetic and trait data could be tackled from the other direction — some lineages may be included in the phylogeny without a corresponding trait value in the dataset — using some sort of data imputation method. A number of recent studies have suggested approaches to accomplish this, some using the parameters of a phylogenetic model (Bruggeman, Heringa & Brandt, 2009; Fagan *et al*., 2013; Guénard, Legendre & Peres-Neto, 2013; Swenson, 2014) and others using a taxonomic sampling model (FitzJohn *et al*., 2014; Sandel *et al*., 2015). These each have their benefits and drawbacks: using phylogenetic models assumes the observed trait values are a random sample of the distribution of trait values, an assumption that may often be egregiously violated (FitzJohn *et al*., 2014), whereas taxonomy-based approaches do not make full use of the structure of the phylogeny and require *ad hoc* assumptions about the sampling distribution for the traits. In any case, all of these involve various assumptions about the unknown states and the validity of these may be difficult to assess in many cases.

The strategies described above are potentially useful for increasing the overlap between the tree and the comparative dataset, but as noted, they may have unintended (and in many cases, poorly understood) consequences for downstream comparative analyses. There is, however, a much simpler approach that has to our knowledge been mostly overlooked by biologists: swap unmatched species in the tree with unmatched species in the data that carry equivalent information content.

Consider a five taxon tree (Figure 2A) of the structure ((((𝒜, 𝔅), 𝒞), 𝒟), ε). If the reconstructed tree contains only taxa 𝒜, 𝒞, 𝒟, and ε, such that the resulting tree has the topology (((𝒜, 𝒞), 𝒟), ε) but our dataset only contains taxa 𝔅, 𝒞, 𝒟, and ε then trait data from taxa 𝔅 can be used in place of trait data for taxa 𝒜 without any loss of information. If we simply dropped unmatched taxa, our analysis would only contain 3 taxa, 𝒞and 𝒟 and ∈, whereas if we exchanged 𝔅 for 𝒜, we would have 4 taxa in our analysis.

**Figure 2:**
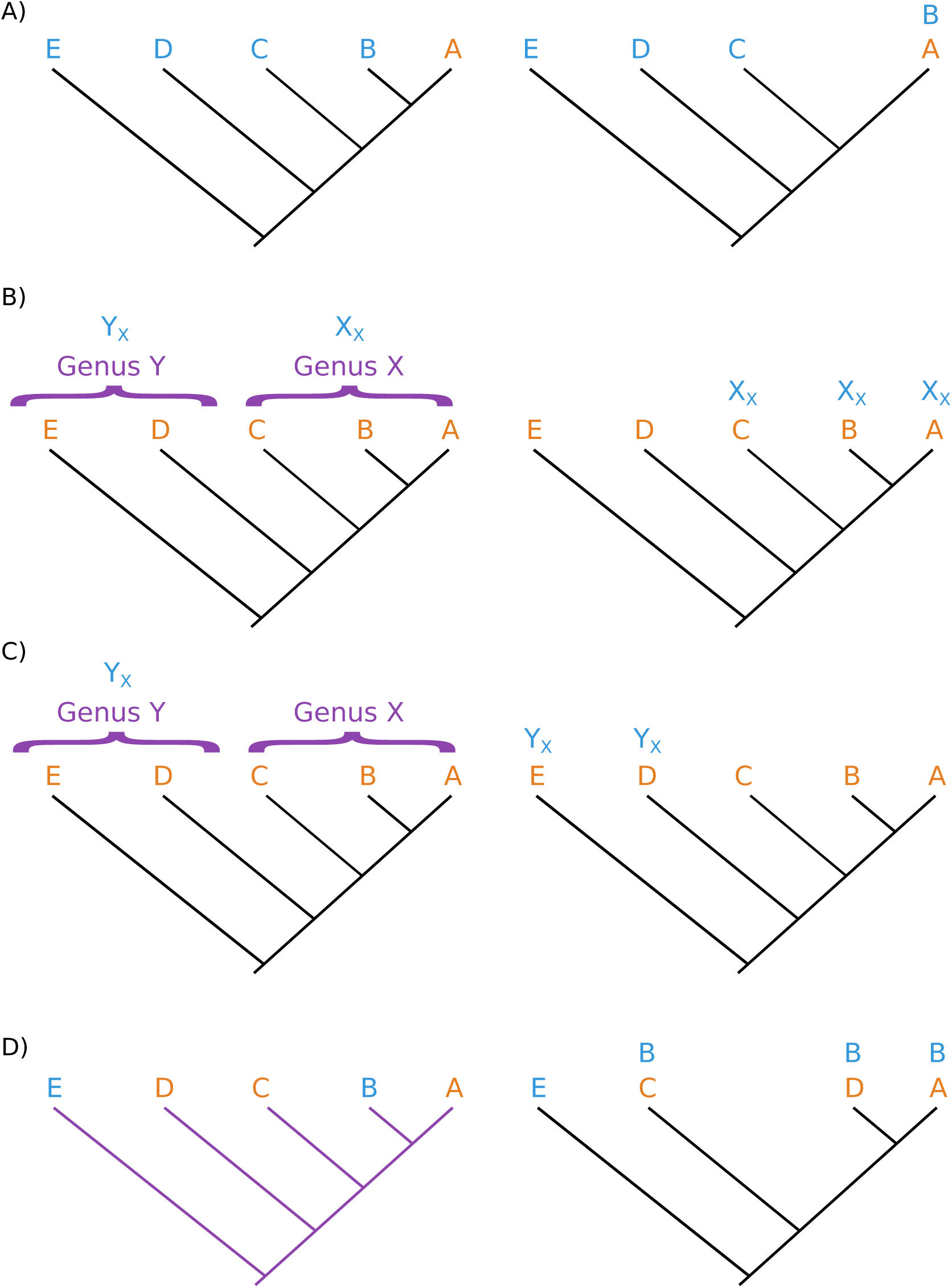
A few examples illustrating the reasoning behind our algorithm. Blue labels indicate species with trait data and orange indicates those without. Panel Because they are sister species, lineages A and B are interchangeable; if we have trait data for one and phylogenetic data for the other, the labels can be swapped (as indicated by the label B over A on the right side). The challenge with incorporating taxonomic information (purple) is that the phylogenetic hypothesis may suggest that named groups are paraphyletic. If the placement of Genus X implies that Genus Y is paraphyletic, then label swaps are only permissible in Genus X if trait data is available for representatives of both X and Y (Panel B). However, if trait data is only available for a representative of genus Y, the label of this lineage can be exchanged with any other member of the genus as all tips from Genus X will be dropped. If one has a topological tree (purple branches; Panel D), a similar principle to the taxonomic case can be applied. Even though lineages C and D are in different positions in the topological tree and the chronogram, because no members of the clade (A,C,D) have data, one can swap in species B for any of these lineages without inducing any splits in the tree.

This trivial example demonstrates that if external knowledge is available, either in the form of a taxonomy or a more comprehensive topological hypothesis, then it is possible to increase the phylogenetic coverage of the data simply swapping phylogenetically equivalent taxa. Of course, simple taxa exchanges such as the above case are logically straightforward and we suspect that this is commonly done in practice by empirical biologists. However, the problem quickly becomes much more complex as the number of mismatches and potential taxa swaps increases, even more so when there is conflict between the supplied topology or taxonomy and the phylogenetic tree being used for analysis. Here we develop a simple algorithm to generate a set of swaps that maximizes the intersection of the phylogenetic tree and comparative data without inducing any new splits in the tree. We have created an efficient implementation of our algorithm which is available as the R package phyndr.

## Taxon-swapping algorithm

Our algorithm takes a time-scaled phylogeny, or *chronogram*, a list of species with *trait data* and an externally supplied *guide* — the guide is distinct from the chronogram. The guide may be either a *topological tree*, a tree containing a more inclusive set of taxa then the chronogram, or else a *taxonomy*. The algorithm differs slightly depending on the type of guide supplied so we deal with these each in turn. We note that technically our algorithm is simply swapping the labels at the tips of the phylogeny but we think it is easier to think of exchanging or swapping taxa, as these are the units of analyses.

We conjecture that whether a topological tree or taxonomy is supplied as a guide, our algorithm will always maximize the intersection of the species in the phylogeny and the dataset without will not inducing any splits that do not occur in the guide (we refer to such swaps as being *permissible*). In this way, our method is conceptually distinct from approaches that randomly place taxa in a tree given some taxonomic knowledge (Kuhn, Mooers & Thomas, 2011; Jetz *et al*., 2012; Thomas *et al*., 2013).

### Using a complete topology

Most modern phylogenetic comparative methods are model-based (see recent reviews by O’Meara, 2012; Pennell & Harmon, 2013), such that branch lengths must be in units of (relative) time for analysis. Using the best estimate of branch lengths is crucial for most modern phylogenetic comparative methods because they infer rates of different evolutionary processes. However, topological information — with no branch length information — may be available from a larger set of the taxa than included in the estimated chronogram: topological trees may come from large supermatrix phylogenies, supertrees, mega-phylogenies (*sensu* Smith, Beaulieu & *Donoghue, 2009*), *or more recently, from synthetic tree alignment graphs (Smith, Brown & Hinchliff, 2013), such as those generated by the Open Tree of Life project (Hinchliff et al.*, 2015). In their raw form, these data sources are not suitable for comparative analyses. However, in combination with a chronogram, phyndr can use this information.

We use a few key definitions to explain the algorithm: all nodes (including tips, nodes without any descendants) can be *complete* or *incomplete*; all descendants of incomplete nodes do not have data, while for complete nodes at least one descendant species has data. This definition follows from the fact that we do not consider any swaps for species that have trait data, even if such swaps are permissible given the topology. Each node in the guide topology is defined by a set of daugther taxa (tip “nodes” are defined by themselves); for each corresponding node in the chronogram, these taxa represent a *candidate set* of possible matches.

We store these at nodes where we might prune the tree down to that node.

The following are the steps in the `phyndr` algorithm:

1. Drop all species from the chronogram that are not in either the data or in the topological tree as these tips are not saveable.
2. Drop species from the topological tree that are not in the data or the chronogram as they are not informative.
3. Flag all tips that have trait data as *complete*, and all other tips and nodes as *incomplete*.
4. Initialise a *candidate set* for each tip and internal node:

a. for tips that have data, the candidate set is the species name;
b. for tips without data, the candidate set is the clade within the topological tree that includes the tip and does not include any other species in the chronogram.
5. In post-order traversal of the chronogram (Felsenstein, 1973), for each node:

a. if any descendant tip/node is complete then this node is complete; the candidate set remains empty;
b. otherwise:

i. compute the descendants of this node within the chronogram;
ii. compute the most recent common ancestor (MRCA) of these descendants in the topological tree;
iii. compute the descendants of that node within the topological tree;
iv. if any descendant in the topological tree is complete, label this node complete;
v. otherwise grow the candidate set to include the descendant nodes’ candidate set, and then clear the descendant nodes’ candidate sets. (This process leaves all species that can be used in exactly one candidate set, and every node will be complete.)
6. Drop all tips in the chronogram with an empty candidate set.

### Using a taxonomic resource

It is likely more common that a taxonomic resource is available for the group of interest. Numerous taxonomic resources are available on the web and emerging tools, such as the R package taxize (Chamberlain & Szöcs, 2013), make it possible to interact with them from within R. For the specific examples in this project we also built a tool, the taxonlookup R package, for building a curated taxonomy of vascular plants (see below for details).

For the taxonomic case, the `phyndr` algorithm works as follows:

Start with a table of taxonomic information; row names are the tip labels in the tree; each column is an increasing taxonomic level (e.g., genus, family, order) that are perfectly nested. Let a *group* be all species at an instance of a taxonomic level (a group may or may not be monophyletic in the chronogram).

For each taxonomic level in decreasing order:

1. Match species in the chronogram to the data; these species are fixed.
2. Drop all species that are in the same *group* as species that have data but which do not have trait data.
3. For each *group* without data, identify if they are monophyletic (i.e., the species in the group form a clade to the exclusion of all other species in the tree).
4. If the *group* contains at least one member with data:

a. if the *group* is monophyletic, collapse into a single tip;
b. otherwise, determine if the group can be *made* monophyletic by dropping other groups that do not have data and if so drop those groups and collapse the focal group.
5. Otherwise (groups with no data), and if the group survived being dropped above:

a. if the group is monophyletic, collapse into a single tip;
b. otherwise leave it alone.

### Dealing with topological conflict

It is important to be explicit about what assumptions we are making when we use a topological tree or taxonomy as a guide. We do not assume that the guide is always correct. Rather, we assume that a group in the topological tree or taxonomy is monophyletic if and only if there is no phylogenetic evidence to contradict this assumption. The phyndr algorithm thus explicitly allows for conflict between the guide and the chronogram. In Figure 2 we walk through some examples of how our algorithm deals with paraphyletic lineages. It is important to keep in mind that monophyly is assessed using species with trait data — even if a lineage renders a group non-monophyletic, it will not affect the permissible swaps if it does map to any trait data. We argue that this set of assumptions is rather weak: it tends toward *not* swapping taxa in cases of conflicting information. And seems likely to be reasonable for many, but not all, stages of development of taxonomic knowledge about specific clades.

### Notes on the algorithm

A number of points are worth considering when applying our algorithm. First, the algorithm does not generate all possible taxon swaps: for lineages that occur in both the tree and the trait data (i.e., those that are considered *complete* in the initial step of the algorithm), we do not consider swaps that exclude the matched species from the final data. If the split (𝒜, 𝓑) exists in the guide (whether topological tree or taxonomy) and both taxa 𝒜 and 𝓑 occur in our data set, but only 𝒜 is in the chronogram, it would be consistent with our algorithm to swap 𝓑 in for 𝓑. However, we have decided to ignore this possibility because it requires making an additional assumption without any gain in information content. (We also note that allowing such swaps would require a more complex algorithm than the one we have proposed.)

Second, while running analyses across multiple permutations of the datasets may be interesting and useful, this does not account for any uncertainty in topology or branch lengths and can therefore not be considered a “posterior distribution” or even a “pseudo-posterior distribution” (*sensu* Thomas *et al*., 2013; Rabosky, 2015). For model-based comparative methods, it is better to consider alternative taxa sets as different realizations of the same process.

Third, our algorithm is restricted to ultrametic phylogenies; taxa are only exchangeable if they are equidistant from their most recent common ancestor, a condition that is only necessarily met when all taxa are sampled at the same time point (see Slater, 2014, for more discussion of this point and its implications for models of trait evolution). So while phylogenetic approaches are becoming increasingly important for analyzing fossil and epidemiological data, alternative strategies will need to be deployed for these cases.

And last, our approach will not be appropriate when testing for trait-dependent diversification (e.g., Maddison, Midford & Otto, 2007; FitzJohn, 2012) or correlations between rates of diversification and rates of trait evolution (e.g., Rabosky *et al*., 2013, 2014) — see Pennell, Harmon & Uyeda 2013 for a discussion of the distinction between these two types of analyses. Dropping tips without any data, which is a step of the `phyndr` algorithm, will tend to push the terminal nodes root-wards and thus bias estimation of diversification rates. Essentially, this is similar to biases introduced by “representative” sampling, in which phylogenies are built using representatives of major taxonomic groups (HÖhna *et al*., 2011; Stadler & Bokma, 2013). The sampling regime introduced by `phyndr` is somewhat different from that studied by FitzJohn, Maddison & Otto (2009), in which all taxa in the phylogeny and assign unknown trait values to species without data, and therefore an alternative correction is needed for this case.

## phyndr R package

We have implemented our algorithm in a new R package `phyndr`. It can be downloaded from the CRAN repository and the development version is available on GitHub (https://github.com/richfitz/phyndr). `phyndr` relies on the ape (Paradis, Claude & Strimmer, 2004) tree structure and diversitree (FitzJohn, 2012) tree manipulation functions. `phyndr` contains two primary functions `phyndr_topology`, and `phyndr_taxonomy` that use topological trees and taxonomies, respectively, as guides. (Generic names can be stripped from taxon labels and used to create a genus-only taxonomy with the function `phyndr_genus`.)

The `phyndr_` functions each generate a `ape::phylo` object with the taxon names relabeled where possible. If multiple relabelings for a given taxa are permissible, one is randomly selected by default but the others are stored in the returned object so that users can generate sets of trees that match to different subsets of the trait data. Note that given the combinatorial nature of the problem, the number of potential relabelings can grow rather quickly and returning the full set may not be possible in R.

## taxonlookup R package

The taxonlookup R package dynamically builds a taxonomic lookup for vascular plants from three canonical sources: The Plant List (The Plant List, 2015), APWeb (Stevens, 2001), and a recently published higher taxa lookup table (Zanne *et al*., 2013, compiled by D.C. Tank, J.M. Eastman, J.M. Beaulieu, W.K. Cornwell, P.F. Stevens, and A.E. Zanne). This will keep the taxonomy resource up-to-date as systematic information improves through time. It will be available on CRAN and on GitHub (https://github.com/wcornwell/taxonlookup); currently only land plants are covered by this resource, but it could be extended to cover other taxa. For taxonlookup we have tried to resolve conflicts and errors in the web-based resources and the package provides output that is can be easily used in conjunction with `phyndr`.

## An example

As a use-case, we applied our algorithm to a recently published time-calibrated phylogeny of vascular plants (Magallón *et al*., 2015) and a database of plant growth form (Zanne *et al*., 2014). The Magallón *et al*. (2015) chronogram contains 798 taxa and many of these were chosen as single representatives of major groups. This feature of the data makes it an ideal situation for applying our algorithm; there will be less opportunity for swapping taxa in datasets where taxon sampling is more phylogenetically clustered. Dropping species for which there was not an exact match between the phylogeny and the trait data would leave us with only 238 taxa — 540 data points are lost!

To improve the overlap, we use `phyndr` in conjuction with the taxonomic table in `taxonlookup` as follows (assuming we have already loaded a `phy` [an `ape::phylo` object] and `dat` [a `data.frame` with rownames equal to the species names] objects into the workspace):

We can get the entire taxonomy table from `taxonlookup` using `plant_lookup`:

~~~
library(taxonlookup)
tax_all <-plant_lookup()
~~~

But for our algorithm, we only need taxonomy for all species in tree and data, which can be obtained with the function `taxonlookup::lookup_table`:

~~~
spp <- unique(c(phy$tip.label, rownames(dat))
tax_spp <- lookup_table(spp, by_species=TRUE)
tax_spp <- tax_spp[,c(“genus”, “family”, “order”)]
~~~

We can then run phyndr to get all permissible taxon swaps:

~~~
library(phyndr)
phyndr_taxonomy(phy, rownames(dat), tax_spp)
~~~

This gives us a comparative data set including 769 species; we have recovered a match for all but 29 of the previously unmatched taxa (Figure 3). (Code to reproduce this example can be found at https://github.com/mwpennell/phyndr-ms.) Similarly, we could have used phyndr_topology and replaced the taxonomic table with a previous topological hypothesis for this group.

**Figure 3:**
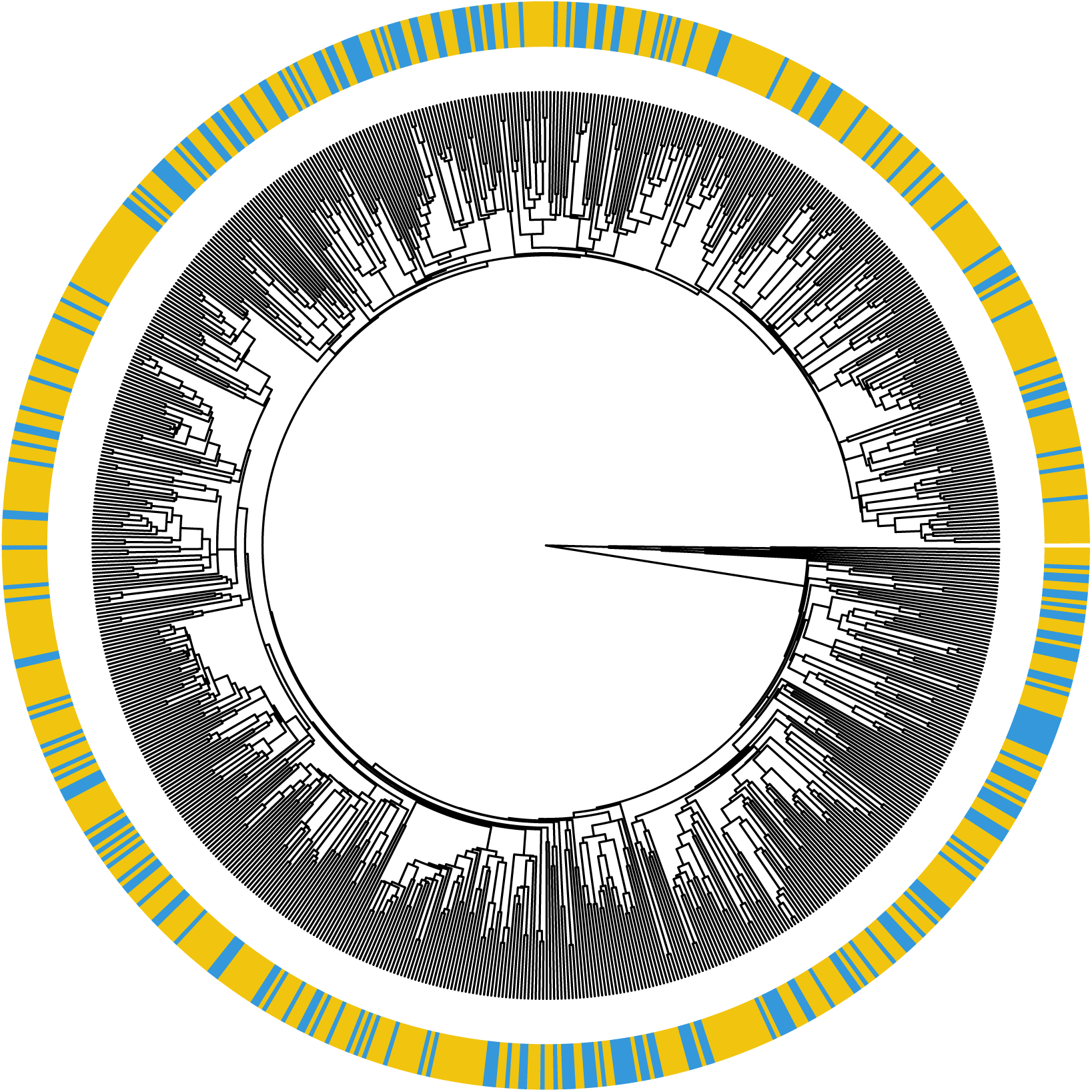
The phylogenetic tree of vascular plants from Magallón *et al*. (2015) after performing label swaps with phyndr using the taxonomic resources in taxonlookup. The original phylogeny contained 798 taxa, only 238 (blue) of which were also in the growth form database. Using phyndr, we were able to find matches for 531 additional taxa (yellow).

## Closing remarks

In recent years, there have been increasing coordination to assemble different species-level data types including observations, traits, genes, and phylogenies (Parr *et al*., 2012). These data sources, while already large and growing, do not overlap completely, and are unlikely to do so for the foreseeable future. As such, any type of synthetic research involving two data sources have a matching problem, and this matching problem will be increasingly common moving into the future. We sought to find a method that maximizes overlap while not introducing a new source of error.

On one level, our method is rather obvious. Simpler versions of the same concept (based on genera only) have been implemented in previous software packages (e.g., phyloGenerator; Pearse & Purvis, 2013). If one is willing to assume that when a node it is present in both the chronogram and a topological or taxonomic hypothesis it can be taken as correct, our method follows from a basic property of ultrametric trees: at any node, the labels of the daughter clades are interchangeable. However, for large, complex topologies with varying degrees of conflict, it is challenging to reason through all permissible label swaps. And even for relatively simple scenarios, automating the process is non-trivial. We believe that phyndr will enable empirical biologists to efficiently and reliably make the most of their data.

A generalised comparative methods workflow consists of the following steps: match exact names; 2) match misspelled and outdated names; 3) substitute close relatives; 4) substitute wherever you can without introducing error; 5) prune the tips with missing data; 6) do analysis. In recent years the tools for some of these steps have improved. For example, step (2) might involve using taxize (Chamberlain & SzÖcs, 2013), Taxonstand (Cayuela *et al*., 2012) and other tools. Our taxonomic package taxonlookup, built upon open datasets from The Plant List (The Plant List, 2015), APWeb (Stevens, 2001), and Open Tree of Life (Hinchliff *et al*., 2015), and the algorithms in phyndr are aimed to maximize effectiveness of steps (3) and (4). We think that our tools will help biologists get the most of their data and be a generally applicable addition to many different comparative methods workflows.

## Acknowledgements

We thank the members of the Tempo and Mode of Plant Trait Evolution working group for stimulating discussions of this problem. In particular, we are grateful to Luke Harmon, Josef Uyeda, Eliot Miller, and Hannah Marx, for advice and comments. We thank Scott Chamberlain and the ROpenSci organization for techincal assistance with the taxonomic resources. This work was supported by the National Evolutionary Synthesis Center (NESCent), NSF #EF- 0905606, Macquarie University Genes to Geoscience Research Centre through the working group. MWP was supported by a NSERC postgraduate fellowship, a NSERC postdoctoral fellowship, and an Izaak Walton Killam Memorial postdoctoral fellowship. This work was also supported by a NSF grant awarded to Luke J. Harmon (DEB 1208912) at the University of Idaho.

## References

Bapst, D.W. (2013) A stochastic rate-calibrated method for time-scaling phylogenies of fossil taxa. Methods in Ecology and Evolution, 4, 724–733.

Bapst, D.W. (2014) Assessing the effect of time-scaling methods on phylogeny-based analyses in the fossil record. Paleobiology, 40, 331–351.

Bruggeman, J., Heringa, J. & Brandt, B.W. (2009) PhyloPars: estimation of missing parameter values using phylogeny. Nucleic acids research, 37, W179–W184.

Cayuela, L., Granzow-de la Cerda, Í., Albuquerque, F.S. & Golicher, D.J. (2012) Taxonstand: An R package for species names standardisation in vegetation databases. Methods in Ecology and Evolution, 3, 1078–1083.

Chamberlain, S.A. & Szöcs, E. (2013) taxize: taxonomic search and retrieval in R. F1000Research, 2.

Davies, T.J., Kraft, N.J.B., Salamin, N. & Wolkovich, E.M. (2012) Incompletely resolved phylogenetic trees inflate estimates of phylogenetic conservatism. Ecology, 93, 242–247.

Fagan, W.F., Pearson, Y.E., Larsen, E.A., Lynch, H.J., Turner, J.B., Staver, H., Noble, A.E., Bewick, S. & Goldberg, E.E. (2013) Phylogenetic prediction of the maximum per capita rate of population growth. Proceedings of the Royal Society B: Biological Sciences, 280, 20130523.

Felsenstein, J. (1973) Maximum-likelihood estimation of evolutionary trees from continuous characters. American journal of human genetics, 25, 471.

FitzJohn, R.G. (2012) Diversitree: comparative phylogenetic analyses of diversification in R. Methods in Ecology and Evolution, 3, 1084–1092.

FitzJohn, R.G., Maddison, W.P. & Otto, S.P. (2009) Estimating trait-dependent speciation and extinction rates from incompletely resolved phylogenies. Systematic biology, 58, 595–611.

FitzJohn, R.G., Pennell, M.W., Zanne, A.E., Stevens, P.F., Tank, D.C. & Cornwell, W.K. (2014) How much of the world is woody? Journal of Ecology, 102, 1266–1272.

Guénard, G., Legendre, P. & Peres-Neto, P. (2013) Phylogenetic eigenvector maps: a framework to model and predict species traits. Methods in Ecology and Evolution, 4, 1120–1131.

Hinchliff, C., Smith, S.A., Allman, J.F., Burleigh, J.G., Chaudhary, R., Cognill, L.M., Crandall, K.A., Deng, J., Drew, B.T., Gazis, R., Gude, K., Hibbett, D.S., Katz, L.A., Laughinghouse IV, H.D., McTavish, E.J., Midford, P.E., Owen, C.L., Ree, R., Rees, J.A., Soltis, D.E., Williams, T. & Cranston, K.A. (2015) Synthesis of phylogeny and taxonomy into a comprehensive tree of life. bioRxiv.

Hinchliff, C.E. & Smith, S.A. (2014) Some limitations of public sequence data for phylogenetic inference (in plants). PLOS ONE, 9, e98986.

HÖhna, S., Stadler, T., Ronquist, F. & Britton, T. (2011) Inferring speciation and extinction rates under different sampling schemes. Molecular biology and evolution, 28, 2577–2589.

Jetz, W., Thomas, G.H., Joy, J.B., Hartmann, K. & Mooers, A.Ø. (2012) The global diversity of birds in space and time. Nature, 491, 444–448.

Jetz, W., Thomas, G.H., Joy, J.B., Redding, D.W., Hartmann, K. & Mooers, A.Ø. (2014) Global distribution and conservation of evolutionary distinctness in birds. Current Biology, 24, 919–930.

Kattge, J., Diaz, S., Lavorel, S., Prentice, I.C., Leadley, P., Bönisch, G., Garnier, E., Westoby, M., Reich, P.B., Wright, I.J. et al. (2011) TRY—a global database of plant traits. Global change biology, 17, 2905–2935.

Kuhn, T.S., Mooers A.Ø. & Thomas, G.H. (2011) A simple polytomy resolver for dated phylogenies. Methods in Ecology and Evolution, 2, 427–436.

Losos, J.B. (1994) An approach to the analysis of comparative data when a phylogeny is unavailable or incomplete. Systematic Biology, 43, 117–123.

Maddison, W.P., Midford, P.E. & Otto, S.P. (2007) Estimating a binary character’s effect on speciation and extinction. Systematic Biology, 56, 701–710.

Magallón, S., Gómez-Acevedo, S., S ánchez-Reyes, L.L. & Hernández-Hernández, T. (2015) A metacalibrated time-tree documents the early rise of flowering plant phylogenetic diversity. New Phytologist.

Martins, E.P. (1996) Conducting phylogenetic comparative studies when the phylogeny is not known. Evolution, 50, 12–22.

O’Meara, B.C. (2012) Evolutionary inferences from phylogenies: a review of methods. Annual Review of Ecology, Evolution, and Systematics, 43, 267–285.

Paradis, E., Claude, J. & Strimmer, K. (2004) APE: analyses of phylogenetics and evolution in R language. Bioinformatics, 20, 289–290.

Parr, C.S., Guralnick, R., Cellinese, N. & Page, R.D. (2012) Evolutionary informatics: unifying knowledge about the diversity of life. Trends in ecology & evolution, 27, 94–103.

Parr, C.S., Wilson, N., Leary, P., Schulz, K.S., Lans, K., Walley, L., Hammock, J.A., Goddard, A., Rice, J., Studer, M. et al. (2014) The encyclopedia of life v2: providing global access to knowledge about life on earth. Biodiversity data journal.

Pearse, W.D. & Purvis, A. (2013) phyloGenerator: an automated phylogeny generation tool for ecologists. Methods in Ecology and Evolution, 4, 692–698.

Pennell, M.W. & Harmon, L.J. (2013) An integrative view of phylogenetic comparative methods: connections to population genetics, community ecology, and paleobiology. Annals of the New York Academy of Sciences, 1289, 90–105.

Pennell, M.W., Harmon, L.J. & Uyeda, J.C. (2013) Is there room for punctuated equilibrium in macroevolution? Trends in Ecology & Evolution, 29, 23–32.

Price, S.A., Hopkins, S.S.B., Smith, K.K. & Roth, V.L. (2012) Tempo of trophic evolution and its impact on mammalian diversification. Proceedings of the National Academy of Sciences, 109, 7008–7012.

Rabosky, D.L. (2015) No substitute for real data: phylogenies from birth-death polytomy resolvers should not be used for many downstream comparative analyses. ArXiv, p. 1503.04978.

Rabosky, D.L., Donnellan, S.C., Grundler, M. & Lovette, I.J. (2014) Analysis and visualization of complex macroevolutionary dynamics: an example from Australian scincid lizards. Systematic biology, 63, 610–627.

Rabosky, D.L., Santini, F., Eastman, J., Smith, S.A., Sidlauskas, B., Chang, J. & Alfaro, M.E. (2013) Rates of speciation and morphological evolution are correlated across the largest vertebrate radiation. Nature communications, 4, 58.

Rolland, J., Jiguet, F., Jønsson, K.A., Condamine, F.L. & Morlon, H. (2014) Settling down of seasonal migrants promotes bird diversification. Proceedings of the Royal Society B: Biological Sciences, 281, 20140473.

Salguero-Gómez, R., Jones, O.R., Archer, C.R., Buckley, Y.M., Che-Castaldo, J., Caswell, H., Hodgson, D., Scheuerlein, A., Conde, D.A., Brinks, E. et al. (2015) The compadre plant matrix database: an open online repository for plant demography. Journal of Ecology, 103, 202–218.

Sandel, B., Gutiérrez, A.G., Reich, P.B., Schrodt, F., Dickie, J. & Kattge, J. (2015) Estimating the missing species bias in plant trait measurements. Journal of Vegetation Science.

Sanderson, M., Donoghue, M.J., Piel, W.H. & Eriksson, T. (1994) TreeBASE: a prototype database of phylogenetic analyses and an interactive tool for browsing the phylogeny of life. American Journal of Botany, 81, 183.

Slater, G.J. (2014) Correction to ‘Phylogenetic evidence for a shift in the mode of mammalian body size evolution at the Cretaceous–Palaeogene boundary’, and a note on fitting macroevolutionary models to comparative paleontological data sets. Methods in Ecology and Evolution, 5, 714–718.

Smith, S.A., Beaulieu, J.M. & Donoghue, M.J. (2009) Mega-phylogeny approach for comparative biology: an alternative to supertree and supermatrix approaches. BMC evolutionary biology, 9, 37.

Smith, S.A., Brown, J.W. & Hinchliff, C.E. (2013) Analyzing and synthesizing phylogenies using tree alignment graphs. PLoS computational biology, 9, e1003223.

Stadler, T. & Bokma, F. (2013) Estimating speciation and extinction rates for phylogenies of higher taxa. Systematic biology, 62, 220–230.

Stevens, P.F. (2001) onwards. angiosperm phylogeny website. Version 12, July 2012, [and more or less continuously updated since].

Swenson, N.G. (2014) Phylogenetic imputation of plant functional trait databases. Ecography, 37, 105–110.

The Plant List (2015) Version 1.1. published on the internet. http://wwwtheplantlistorg/, accessed 2 May.

Thomas, G.H., Hartmann, K., Jetz, W., Joy, J.B., Mimoto, A. & Mooers, A.Ø. (2013) PASTIS: an R package to facilitate phylogenetic assembly with soft taxonomic inferences. Methods in Ecology and Evolution, 4, 1011–1017.

Vellend, M., Cornwell, W.K., Magnuson-Ford, K. & Mooers, A.Ø. (2011) Measuring phylogenetic biodiversity. Biological diversity: frontiers in measurement and assessment Oxford University Press, Oxford, UK, pp. 194–207.

Wheeler, D.L., Barrett, T., Benson, D.A., Bryant, S.H., Canese, K., Chetvernin, V., Church, D.M., DiCuccio, M., Edgar, R., Federhen, S. et al. (2007) Database resources of the national center for biotechnology information. Nucleic acids research, 35, D5–D12.

Zanne, A.E., Tank, D.C., Cornwell, W.K., Eastman, J.M., Smith, S.A., FitzJohn, R.G., McGlinn, D.J., O’Meara, B.C., Moles, A.T., Reich, P.B., Royer, D.L., Soltis, D.E., Stevens, P.F., Westoby, M., Wright, I.J., Aarssen, L., Bertin, R.I., Calaminus, A., Govaerts, R., Hemmings, F., Leishman, M.R., Oleksyn, J., Solits, P.S., Swenson, N.G., Warman, L. & Beaulieu, J.M. (2014) Three keys to the radiation of angiosperms into freezing environments. Nature, 506, 89–92.

Zanne, A.E., Tank, D.C., Cornwell, W.K., Eastman, J.M., Smith, S.A., FitzJohn, R.G., McGlinn, D.J., O’Meara, B.C., Moles, A.T., Reich, P.B., Royer, D.L., Soltis, D.E., Stevens, P.F., Westoby, M., Wright, I.J., Aarssen, L., Bertin, R.I., Calaminus, A., Govaerts, R., Hemmings, F., Leishman, M.R., Oleksyn, J., Solits, P.S., Swen-son, N.G., Warman, L. & Beaulieu, J.M. (2013) Data from: Three keys to the radiation of angiosperms into freezing environments. Dryad Digital Repository. doi:10.5061/dryad.63q27.2.

